# The formation of ubiquitin rich condensates triggers recruitment of the ATG9A lipid transfer complex to initiate basal autophagy

**DOI:** 10.1101/2023.11.28.569058

**Authors:** DG Broadbent, CM McEwan, TM Tsang, DM Poole, BC Naylor, JC Price, JC Schmidt, JL Andersen

## Abstract

Autophagy is an essential cellular recycling process that maintains protein and organelle homeostasis. ATG9A vesicle recruitment is a critical early step in autophagy to initiate autophagosome biogenesis. The mechanisms of ATG9A vesicle recruitment are best understood in the context of starvation-induced non-selective autophagy, whereas less is known about the signals driving ATG9A vesicle recruitment to autophagy initiation sites in the absence of nutrient stress. Here we demonstrate that loss of ATG9A or the lipid transfer protein ATG2 leads to the accumulation of phosphorylated p62 aggregates in the context of basal autophagy. Furthermore, we show that p62 degradation requires the lipid scramblase activity of ATG9A. Lastly, we present evidence that poly-ubiquitin is an essential signal that recruits ATG9A and mediates autophagy foci assembly in nutrient replete cells. Together, our data support a ubiquitin-driven model of ATG9A recruitment and autophagosome formation during basal autophagy.

## Introduction

The recycling of cellular material through macroautophagy (referred to here as autophagy) maintains organelle and protein homeostasis. Defects in autophagy are associated with numerous human diseases, including metabolic disorders, neurodegeneration, infectious disease, and autoimmunity (Mizushima and Levine, 2020). Autophagy is characterized by the formation of a double membrane vesicle called the autophagosome, which engulfs cellular material and ultimately fuses with the lysosome to degrade and recycle its contents (reviewed in (Debnath, Gammoh and Ryan, 2023)). A variety of proteins regulate autophagosome growth, closure, and fusion with the lysosome. Of the >40 core autophagy-regulating proteins identified, approximately 17 are considered essential and carry the ‘ATG’ designation. These ATG proteins are evolutionarily conserved and required for non-selective and selective forms of autophagy (Vargas et al., 2023). One essential yet still incompletely understood protein that acts at the earliest steps of autophagy is ATG9A (Karanasios et al., 2016; Yamamoto et al., 2012). Loss of ATG9A causes a severe autophagy-deficient phenotype in cultured cells and homozygous deletion of ATG9A in mice is lethal (Kuma et al., 2004).

ATG9A is the only multi-pass transmembrane protein among the core ATG proteins. Under nutrient replete conditions, ATG9A associates with small lipid vesicles that move through the ER, Trans-Golgi Network, and endomembrane vesicle systems (Imai et al., 2016; Mari et al., 2010; Nishimura et al., 2017; Orsi et al., 2012; Young et al., 2006). In response to an autophagy stimulus like starvation, ATG9A vesicles help recruit other ATG proteins and ultimately expand into phagophores (Judith et al., 2019). Recent work suggests that ATG9A acts as a lipid scramblase, receiving phospholipids from the lipid transferase ATG2 and distributing these lipids between the inner and outer leaflets of the growing phagophore membrane, thereby regulating autophagosome membrane curvature and lipid composition, which is essential for autophagy progression (Maeda et al., 2020; Matoba et al., 2020).

In contrast to starvation-induced autophagy, less is known about mechanisms of autophagy in the absence of nutrient stress, which we refer to here as basal autophagy—a broad term that encompasses several mechanistically different forms of autophagy that occur in nutrient-replete conditions. In general, basal autophagy is cargo specific. For example, a form of basal autophagy called aggrephagy performs the pro-survival function of pruning toxic protein aggregates from the cell (reviewed in (Adriaenssens, Ferrari and Martens, 2022)). As such, defects in aggrephagy are associated with proteinopathy diseases, including Parkinson’s disease and amyotrophic lateral sclerosis (ALS) (Mizushima and Levine, 2020). The specificity of basal autophagy is driven at least in part by autophagy adapter proteins, including p62, TAX1BP1, NBR1, and OPTN, which interact with poly-ubiquitinated substrates and gather them for engulfment by the autophagosome (reviewed in (Adriaenssens, Ferrari and Martens, 2022)). For example, through its ubiquitin association (UBA) domain, p62 binds to poly-ubiquitinated misfolded proteins and oligomerizes to form p62- and ubiquitin-rich phase separated droplets, referred to here as ubiquitin-rich condensates, which can be engulfed by autophagosomes (Bjorkoy et al., 2005; Pankiv et al., 2007; Turco et al., 2021; Wurzer et al., 2015; Zaffagnini et al., 2018). Other autophagy adapters are also recruited to these ubiquitin-rich condensates, including NBR1 and TAX1BP1, which likely help package the condensate for engulfment by the autophagosome (Turco et al., 2021). Therefore, in this model, the formation of a ubiquitin-rich condensate seems to be an initial trigger to recruit autophagy machinery, which, in turn, generates the autophagosome in a cargo-directed manner. Indeed, cell treatments that cause a build-up of ubiquitin-rich condensates trigger a corresponding accumulation of ATG9A at the condensates, suggesting that ATG9A is somehow recruited early to these sites of selective autophagy even in the absence of starvation-induced signaling (Kannangara et al., 2021). However, the mechanism of ATG9A recruitment to these sites is not completely understood.

In this study, we used proteomics coupled with D_2_O labeling to measure the impact of ATG9A deletion on proteome flux in nutrient replete cells. Our results show that loss of ATG9A affects the degradation rates of many proteins, but none more than the autophagy adapter p62. We demonstrate that the loss of ATG9A promotes the accumulation of phosphorylated forms of p62 in large cytoplasmic clusters consistent with ubiquitin-rich condensates. Deletion of other core ATG proteins (ATG13 and ATG5) does not entirely phenocopy ATG9A deletion in regard to p62 phosphorylation and condensate size. However, deletion of ATG2, a partner of ATG9A in mediating lipid transfer and phagophore expansion, phenocopies ATG9A loss, supporting the critical role of this lipid transfer complex in basal autophagy. We also show that p62 flux requires ATG9A lipid scramblase activity and deletion of ATG9A abrogates ATG2A recruitment to ubiquitin-rich condensates. Lastly, we show that poly-ubiquitin is required and sufficient to recruit ATG9A to p62-positive condensates. Taken together, our data support a model in which the accumulation of poly-ubiquitin in clusters or condensates is a trigger to recruit ATG9A and initiate their degradation in the context of basal autophagy.

## Results

### Quantitative proteome-level measurement of protein turnover identifies p62 as an ATG9A-dependent target in basal autophagy

To determine how loss of ATG9A affects protein flux across the proteome during fed conditions, we used a quantitative deuterium labeling LC-MS/MS approach (Mathis et al., 2017; Naylor et al., 2017) to derive synthesis and degradation rates for endogenous proteins in HEK293T wild-type (WT) and HEK293T ATG9A knock-out (KO) cells (Fig. 1A). Of the 4,025 proteins identified, 517 of these were considered to have sufficient quality and quantity within all replicates to be considered for analysis of turnover rates (Fig. 1B). As we expected, the largest fraction of the 517 proteins (208) encompassed proteins that showed decreased degradation rates in ATG9A KO cells. A GO term and Pathway analysis of these proteins revealed an enrichment in vesicle trafficking pathway and metabolism (Figure 1C). We found that the autophagy adapter p62 had the most significantly decreased degradation rate in ATG9A KO cells of all proteins detected (Figure 1B and D). This strong effect of ATG9A KO on p62 stability is also corroborated by previous CRISPR screens (DeJesus et al., 2016; Shoemaker et al., 2019). Taken together, our proteome flux measurements suggest that ATG9A-dependent basal autophagy targets cytosolic vesicle trafficking and metabolism proteins, and most prominently the autophagy adapter p62 and its associated cargo.

**Figure 1.**
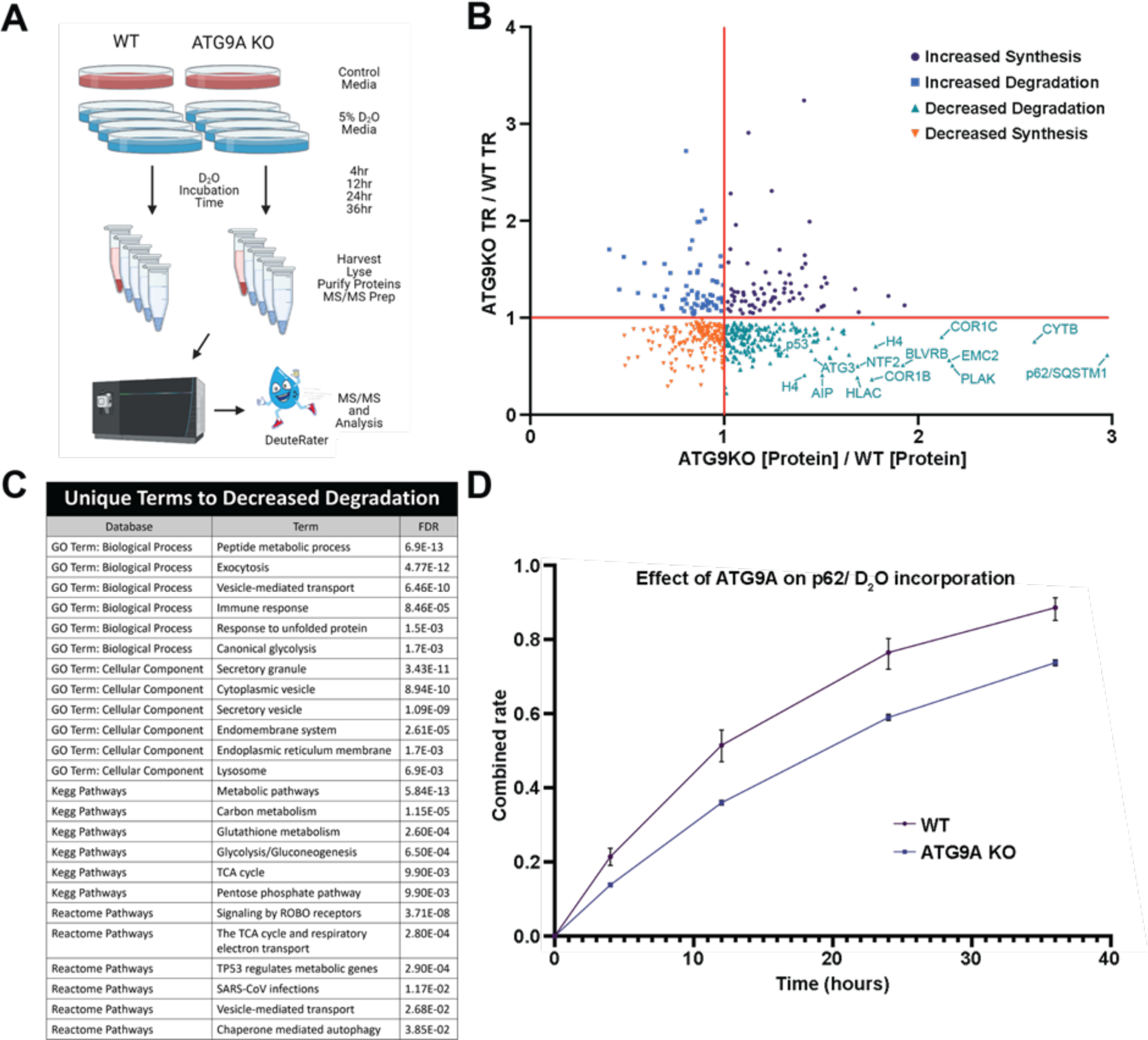
Quantitative proteome turnover analysis comparing WT and ATG9A KO cells. **(A)** Schematic showing the workflow using label-free quantitation (control media, n=3) and metabolic deuterium incorporation (D2O media, n=3 collected @ 4, 12, 24, and 36 hr) to derive protein abundance and turnover rates (TR) in WT and ATG9A KO HEK293T cells. **(B)** Concurrent changes in abundance and turnover, shown as fold differences (ATG9A KO vs WT cells). The values were derived from a biological triplicate of the experiment in panel A with error bars omitted for clarity of presentation **(C)** Ontologies enriched in the Decreased Degradation quadrant (positive [Protein] with negative turnover) for ATG9AKO vs WT as scored by Panther. **(D)** Metabolic labeling of P62, as an example of the protein turnover data. Error bars represent SD between technical replicates (n=3) for the fraction of new P62 detected by mass spectrometry at the indicated time.

### Phosphorylated p62 accumulates prominently in cells lacking the ATG9A-ATG2 lipid transfer complex

To determine if the loss of other key autophagy proteins would affect p62 degradation similarly to loss of ATG9A, we knocked out ATG5 and ATG13 in HEK293T cells to generate LC3 lipidation-deficient (ATG5 KO) and cells defective in the formation of the ULK1 complex (ATG13 KO). Quantification of total p62 levels and flux revealed little difference between the ATG9A KO, ATG5 KO, and ATG13 KO lines. However, the ATG9A KO line displayed a prominently upshifted p62 band compared to the other KO lines. (Fig. 2A-B). Using phosphospecific-antibodies to detect known phosphorylation sites on p62, including pS269, pS349, and pS403, we confirmed that the upshifted p62 band in ATG9 KO cells is phosphorylated at these sites (Fig. 2C-E). Bafilomcycin treatment had little effect on the turnover of phospho- and unmodified p62, suggesting loss of ATG9A resulted in the accumulation of a phosphorylated form of p62 that is stalled in the autophagy process (Fig. 2C-E).

**Figure 2.**
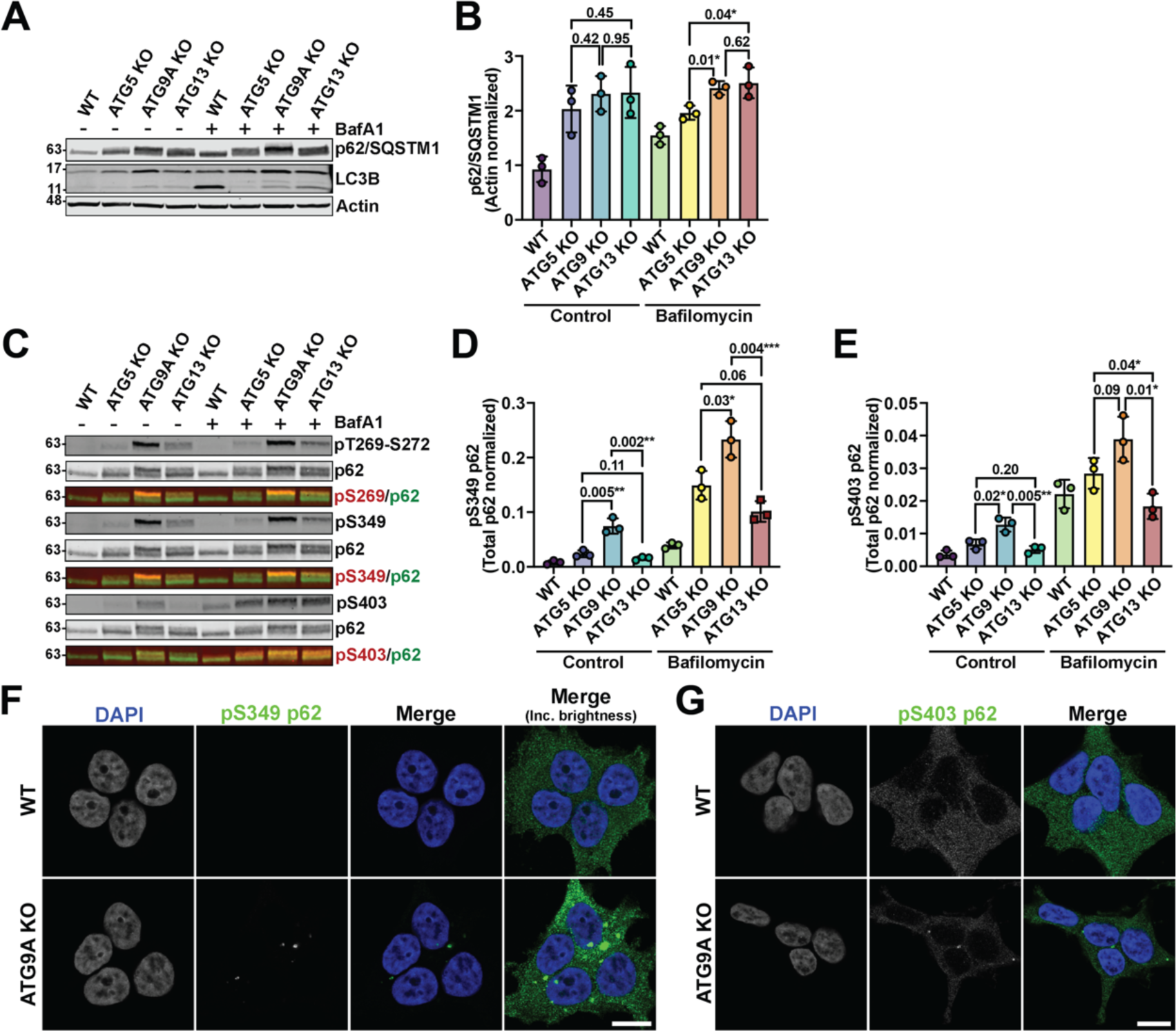
Phosphorylated p62 accumulates prominently in ATG9A KO HEK293T cells. **(A)** Western blot demonstrating the defects of p62 degradation in ATG5, ATG9A, and ATG13 KO cell lines. **(B)** Quantification of p62 immunoblot signal normalized to actin from 3 replicates. Error bars represent SD. *P*-values were calculated using a two-tailed student *t*-test. **(C)** Immunoblot showing the buildup of phospho-p62 in the indicated ATG protein KO cell lines. **(D-E)** Quantification (as done in panels B-C) of pS349 and pS403 on p62 from 3 biological replicates of immunoblots as shown in panel D. Error bars represent SD. *P*-values were calculated using a two-tailed student *t*-test. **(F-G)** Images depicting the accumulation of pS349 or pS403 p62 in WT and ATG9A KO HEK293T cells (scale bars = 10 µm).

Previous work demonstrated that phosphorylation of p62 at S349 and S403 increased p62 affinity to cargo and/or poly-ubiquitin (Ichimura et al., 2013; Matsumoto et al., 2015; Matsumoto et al., 2011; Pilli et al., 2012). This suggests these phosphorylation sites mark the pool of p62 at ubiquitin-rich condensates. Supporting this idea, we found that the S349- and S403-phosphorylated forms of p62 were concentrated within large cytosolic droplets consistent in morphology with p62-positive ubiquitin-rich condensates (Fig. 2F-G) (Kannangara et al., 2021). We also observed the autophagy adapter TAX1BP1 at these condensates (Fig. S1), supporting the idea that these are sites of autophagy initiation in fed cells.

Recent structural and modeling work has suggested that ATG9A is a lipid scramblase that forms a complex with the lipid transfer proteins ATG2A and ATG2B to initiate autophagosome formation (Ghanbarpour et al., 2021; Maeda et al., 2020; Matoba et al., 2020; van Vliet et al., 2022). Together, this ATG9A-ATG2 complex is proposed to maintain lipid balance across the autophagosome membrane as it expands. Therefore, given their connected roles, we hypothesized that loss of ATG2A/B would mimic loss of ATG9A and also lead to the accumulation of phosphorylated p62. To test this hypothesis, we used a recently established cell in which the HaloTag is homozygously inserted at the N-terminus of genomic *atg9a* locus in U2OS cells (Broadbent et al., 2023). HaloTagged ATG9A is fully functional and supports LC3-lipidation and p62 degradation (Broadbent et al., 2023). In the context of this HaloTag-ATG9A U2OS cell line we knocked out ATG2A/B, ATG5, or ATG9A (Broadbent et al., 2023) and confirmed that p62 degradation and phosphorylation showed a similar dependency on ATG9A in these cells (Fig. 3A-B). Deletion of ATG2A/B phenocopied loss of ATG9A resulting in an accumulation of phosphorylated p62 (Fig. 3C-F). We also attempted to determine whether deletion of the ATG9A paralog ATG9B in our ATG9A KO cells would cause an even more severe defect in p62 accumulation. However, we were unable to recover any surviving ATG9A/B double knock-out cells. Nevertheless, our data suggested that loss of the ATG9A-ATG2 lipid transfer complex causes a significant accumulation of phosphorylated p62 (Fig. 3C-F). To determine whether ATG9A scramblase function is required for degradation of the phosphorylated pool of p62, we expressed Halo-ATG9A WT or an ATG9A mutant with reduced lipid scramblase activity (ΔScramblase or M33; K321L, R322L, E323L, T419W (Maeda et al., 2020)) in ATG9A KO U2OS cells using a inducible expression cassette. After 24 hours of doxycycline induced ATG9A expression, we found that ATG9A-M33 failed to rescue p62 degradation in the ATG9A KO line (Fig. 3G-H). Together, these results suggest that deletion of the ATG9A-ATG2 lipid transfer complex causes a large accumulation of phosphorylated p62 that is not observed when ATG13 or ATG5 are knocked out.

**Figure 3.**
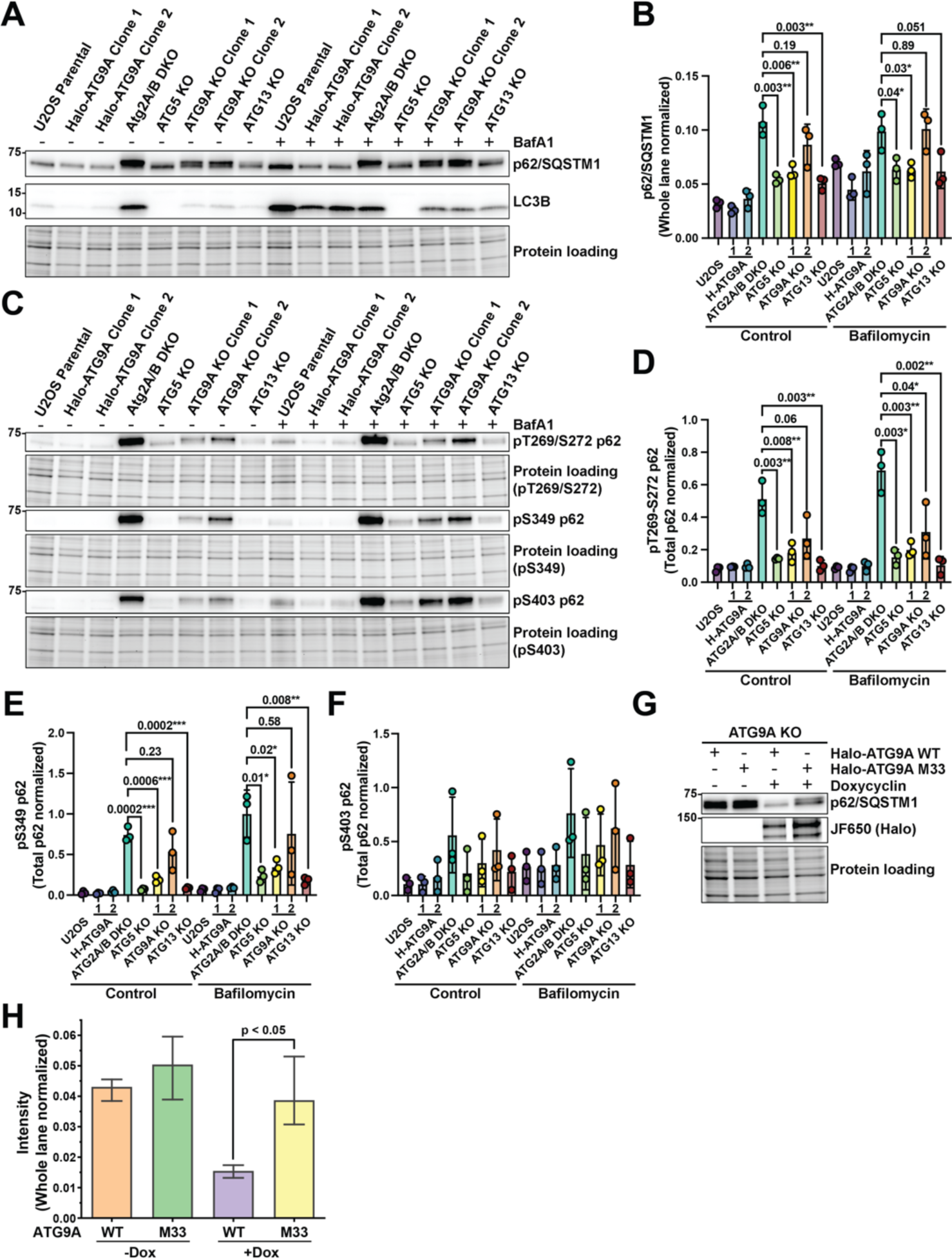
The ATG9A-ATG2 lipid transfer complex is required for degradation of phosphorylated p62. **(A)** Western Blot demonstrating the defects of p62 degradation in ATG5, ATG9A, and ATG13 KO U2OS cell lines. **(B)** Quantification of the triplicate western blots in (A). Error bars represent SD. *P*-values were calculated using a two-tailed student *t*-test. **(C)** Western blot showing the accumulation of phospho-p62 in the indicated ATG KO U2OS cell lines. **(D-F)** Quantification of the phospho-p62 western blots in (C). Error bars represent SD. *P*-values were calculated using a two-tailed student *t*-test. **(G)** Fluorescent gel and western blot showing the degradation of p62 in cells stably expressing WT and M33 Halo-ATG9A variants in the absence and presence of doxycycline. **(H)** Quantification of triplicate western blots represented in (G). Error bars represent SD. *P*-values were calculated using a two-tailed student *t*-test.

### The formation of ubiquitin-rich condensates triggers the recruitment of ATG9A to initiate autophagy

In previous work, we have demonstrated that perturbations that increase ubiquitin-rich condensate size result in an accumulation of ATG9A at these condensates, which also co-localize with p62, TAX1BP1 and other autophagy markers (Kannangara et al., 2021). However, the molecular mechanism of ATG9A recruitment to these ubiquitin-rich condensates remains unclear. Notably, the ULK1 complex scaffold FIP200, which tethers the ULK1 complex to p62 and other adapters at ubiquitin-rich condensates (Ravenhill et al., 2019; Turco et al., 2019; Vargas et al., 2019), is not required for ATG9A recruitment to these aggregates (Fig. S2), suggesting that other components of the condensate recruit ATG9A. ATG9A is observed at the earliest steps of autophagosome growth (Ren et al., 2023; Sawa-Makarska et al., 2020; Yamamoto et al., 2012), which suggests at least two models of autophagy initiation by ATG9A: the clustering of ATG9A vesicles may occur first to generate nascent autophagosomal membrane to which p62, OPTN, TAX1BP1 and other adapters recruit cargo and downstream autophagy machinery, or alternatively, the ubiquitin-rich condensates form first to create a platform for ATG9A recruitment. The latter model is supported by evidence that OPTN is required for ATG9A recruitment to artificially induced ubiquitin condensates (Yamano et al., 2020). To distinguish between these two models of ATG9A-mediated basal autophagy, we took advantage of HCT-116 cells in which we appended an HA tag in-frame to the C-terminus of the *atg9a* gene, which allows us to track endogenous ATG9A accumulation at ubiquitin-rich condensates (Kannangara et al., 2021). Treatment of WT cells with wortmannin, which blocks autophagosome growth, resulted in a minor accumulation of small ubiquitin-rich foci positive for p62 and ATG9A (Fig. 4A-C). However, deletion of p62, the scaffold of ubiquitin condensate formation (Ciuffa et al., 2015; Turco et al., 2021), eliminated the ATG9A-positive aggregates (Fig. 4A-C). To define the relationship between p62 condensate formation and ATG9A accumulation, we deleted ATG13 in these cells, which led to ATG9A accumulation at very large ubiquitin-rich condensates. Deletion of p62 eliminated these large condensates, and reverted ATG9A back to a normal cellular distribution (Fig. 4A-C). This suggests that the formation of ubiquitin-rich condensates precedes ATG9A recruitment.

**Figure 4.**
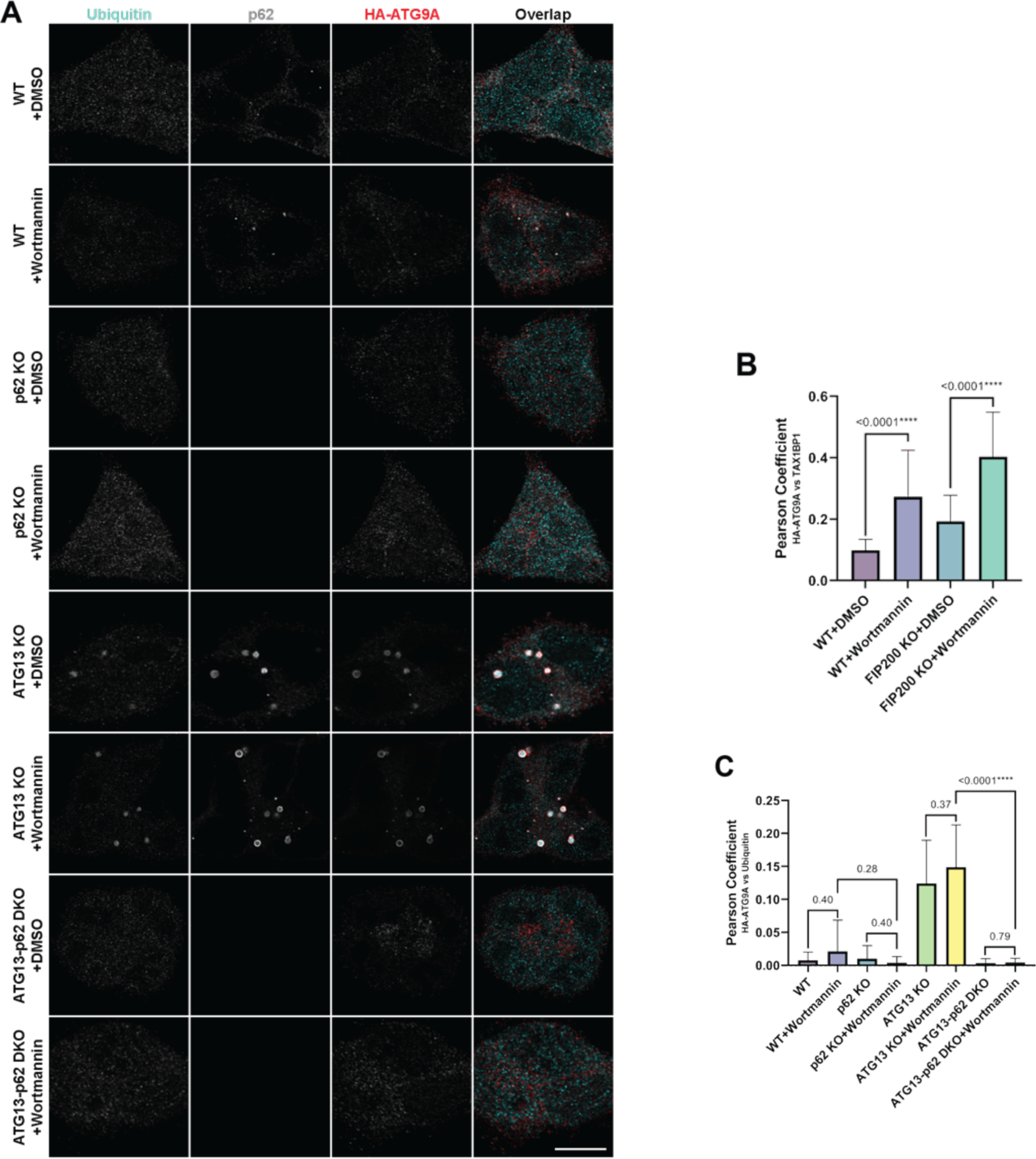
p62-mediated formation of ubiquitin-rich condensates is required for ATG9A recruitment. **(A)** Confocal imaging of endogenous p62, ubiquitin and endogenously HA-tagged ATG9A in WT, ATG13 KO and ATG13-p62 double KO (dKO) HCT-116 cells (scale bars = 10 µm). Where indicated, cells were treated for 4 hours with 1 uM wortmannin. (B) Quantification of ATG9A-HA and TAX1BP1 colocalization by Pearson’s coefficient Quantitation is from 3 replicates with error bars representing SD. *P*-values were calculated using a two-tailed student *t*-test for pair-wise comparison. (C) Quantification of ATG9A-HA and ubiquitin colocalization from 3 biological replicates with error bars representing SD. *P*-values were calculated using a two-tailed student *t*-test as in panel B.

We recently developed a panel of endogenously tagged autophagy proteins to quantitatively measure the formation of autophagy foci by live cell imaging (Barnaba et al., 2023; Broadbent et al., 2023). Our experiments with these cell lines suggest that the autophagy machinery rapidly assembles and disassembles into foci, but only a fraction of these foci ultimately mature into LC3 containing autophagosomes (Broadbent et al., 2023). A growing body of literature suggests that poly-ubiquitin chains attached to misfolded or aggregated proteins are the trigger that nucleates autophagosome formation at ubiquitin-rich condensates (Ravenhill et al., 2019; Sun et al., 2018; Turco et al., 2021; Turco et al., 2019; Zaffagnini et al., 2018). We therefore hypothesized in the context of basal autophagy in nutrient rich media, autophagy factors can form persistent foci when they are tethered to a degradation target, for instance poly-ubiquitin at ubiquitin-rich condensates. To determine whether the formation of poly-ubiquitinated aggregates triggers foci formation of autophagy factors, we transiently inhibited ubiquitination, which should reduce the formation of new ubiquitin-rich condensates. In this experiment, we used Halo-ATG13 as a readily detectable marker of autophagy foci formation while also tracking the colocalization of ATG13 with LC3 to assess autophagosome maturation. Inhibition of poly-ubiquitination with the E1 ubiquitin-activating enzyme inhibitor MLN4924 (MLN) reduced the formation of ATG13 foci that colocalized with LC3B. The inhibition of poly-ubiquitination also reduced ATG13 foci that were not colocalized yet with LC3B, suggesting a role for poly-ubiquitin in the initial formation of autophagy foci (Fig. 5A; Videos 1 and 2). In addition, we analyzed the lifetime of ATG13 foci, which was largely unchanged in control and MLN treated cells, perhaps reflecting ATG13 foci that form at already established (prior to MLN treatment) ubiquitin-rich condensates—and once formed, they persist as usual (Fig. 5B; Videos 1 and 2). Furthermore, in both control and MLN treated cells, the lifetime of ATG13 foci that colocalized with LC3 was significantly longer than ATG13 that did not colocalize with LC3 signal, consistent with the idea that LC3-positive foci have passed a commitment step to form autophagosomes (Fig. 5B; Videos 1 and 2) (Broadbent et al., 2023). Taken together, these observations suggest that the formation of ubiquitin-rich condensates/aggregates creates a platform for ATG protein recruitment and autophagosome initiation.

**Figure 5.**
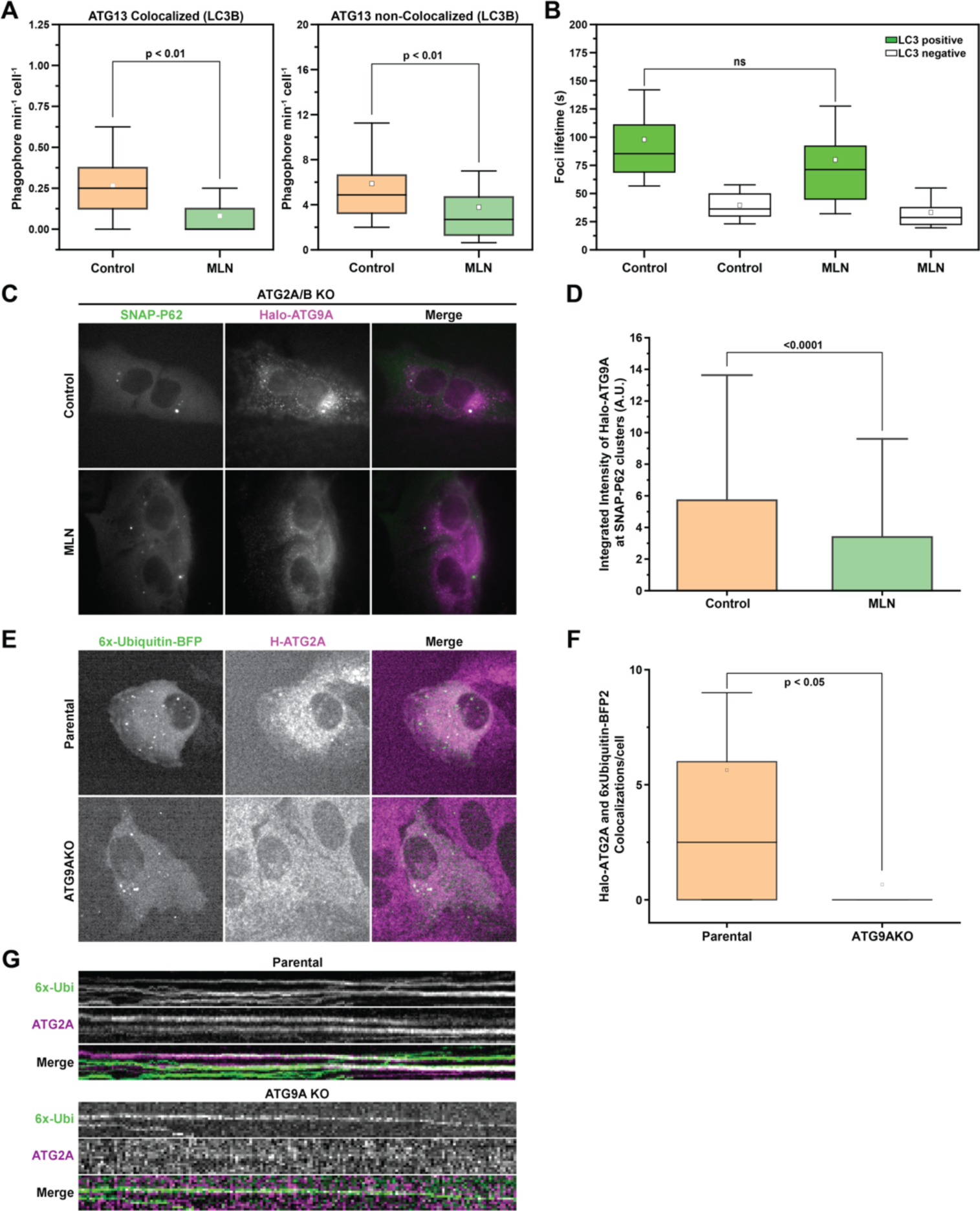
Poly-ubiquitin is required and sufficient for recruitment of the ATG9A-ATG2A lipid transfer complex. **(A)** Rates of ATG13 foci formation divided into Colocalized and non-Colocalized foci in U2OS cells endogenously expressing Halo-ATG13 and stably expressing GFP-LC3B treated with DMSO or MLN. *P*-values were calculated using a two-tailed student *t*-test. **(B)** ATG13 foci length distribution from the data shown in (A). *P*-values were calculated using a two-tailed student *t*-test. **(C)** Example images showing the difference in recruitment of endogenous Halo-ATG9A to SNAP-p62 aggregates in the presence and absence of MLN. **(D)** Quantification of the amount of Halo-ATG9A that is recruited to SNAP-P62 aggregates shown in (C). *P*-values were calculated using a two-tailed student *t*-test. **(E)** Representative images showing the recruitment of Halo-ATG2A to 6x-Ubiquitin-BFP2 aggregates in WT and ATG9AKO cells. **(F)** Counts of colocalized foci shown in (E). **(G)** kymograph showing representative foci of Halo-ATG2A and 6x-ubiqutin colocalization over-time from the experiment shown in (E). *P*-values were calculated using a two-tailed student *t*-test.

Next, we asked whether the inhibition of ubiquitination prevented ATG9A recruitment to p62-positive structures. To visualize robust recruitment of ATG9A to large condensates, we used ATG2A/B knock out cells in which ATG9A vesicles are recruited to autophagy substrates that fail to be degraded (Olivas et al., 2023). To visualize ATG9A at these condensates, we used our endogenously tagged Halo-ATG9A cell line and expressed SNAP-p62 by stable integration into the AAVS1 locus. To distinguish between ATG9A foci that were already present prior to MLN treatment and ATG9A recruited to nascent condensates during MLN treatment, we blocked existing Halo-ATG9A with nonfluorescent HaloTag ligand, followed by incubation with the fluorescent JF646 HaloTag ligand. Using this approach, we only visualized newly synthesized Halo-ATG9A rather than Halo-ATG9A that may already be accumulated at pre-existing condensates. When ubiquitination was inhibited with MLN, we observed reduced recruitment of ATG9A to p62 condensates (Fig. 5C-D; Videos 3 and 4), suggesting that poly-ubiquitination is required for ATG9A recruitment to these condensates.

To test this model further, we determined whether the clustering of poly-ubiquitin is sufficient to recruit ATG9A. However, our previous work demonstrated that ATG9A does not detectably accumulate at autophagy foci in cells with an intact autophagy pathway (Broadbent et al., 2023). To overcome this challenge, we used two different approaches. First, we transiently overexpressed SNAP-tagged ATG9A and 6x-ubiquitin-BFP2 to form artificial ubiquitin-rich condensates. In this experiment, we only observed detectable accumulation of SNAP-ATG9A at 6x-ubiquitin-BFP2 clusters in cells also co-overexpressing OPTN (Fig. S3), which corroborates the observation by Matsuda and colleagues that OPTN promotes ATG9A localization to artificial ubiquitin clusters (Yamano et al., 2020). We also tested the possibility that poly-ubiquitination of ATG9A itself may be required for recruitment to ubiquitin-rich condensates. However, lysine-to-arginine mutations at ATG9A ubiquitination sites (K581 and K838) that were reported as necessary for formation of VPS34-UVRAG complexes (Wang et al., 2022) did not affect ATG9A recruitment to the artificial ubiquitin clusters (Fig. S3). Lastly, we used our endogenously tagged Halo-ATG2A cell line, which detectably accumulates at phagophores, as a marker for the formation of the ATG9A-ATG2 lipid transfer complex. We observed robust ATG2A recruitment to 6x-ubiquitin-BFP2 condensates (Fig. 5E-G; Videos 5 and 6). Importantly, the recruitment of ATG2 to 6x-ubiquitin-BFP2 clusters depended on ATG9A (Fig. 5E-G; Videos 5 and 6). Taken together, these observations suggest that the formation poly-ubiquitin condensates, likely together with autophagy adapters like OPTN, recruits the ATG9A-ATG2 lipid transfer complex to initiate autophagosome growth in the context of basal autophagy.

## Discussion

The work presented in this study, together with a growing body of published literature (Turco et al., 2021; Turco et al., 2019; Vargas et al., 2019; Wurzer et al., 2015; Zachari et al., 2019; Zaffagnini et al., 2018), suggest that the initiation of autophagy in the absence of acute nutrient stress occurs as ubiquitin-binding adapters gather ubiquitinated cargo (e.g. misfolded proteins) to form ubiquitin-rich condensates, which provide a platform to recruit ATG9A and begin assembling the ATG9A-ATG2 lipid transfer complex to initiate autophagosome biogenesis. However, important questions remain. It is still not clear what initially recruits and tethers ATG9A vesicles to the ubiquitin-rich condensate. Because poly-ubiquitin is a central feature of the diverse substrates of basal autophagy, it seems likely that ubiquitin itself, perhaps cooperatively with autophagy adapters like OPTN (Yamano et al., 2020), interacts directly with ATG9A or another component of the ATG9A vesicle. Our work supports this model but does not entirely rule out other possibilities. For example, it is possible that other core autophagy proteins localize to the condensate upstream of ATG9 and facilitate its recruitment. FIP200 was a potential candidate, but our results demonstrate that FIP200 is not necessary for ATG9A recruitment to P62-condensates. Recent structural and molecular studies also demonstrated an interaction between the C-terminus of ATG9A and the ATG13:ATG101 dimer (Kannangara et al., 2021; Ren et al., 2023). ATG13, like FIP200, is dispensable for ATG9A recruitment to ubiquitin-rich condensates in fed conditions (Kannangara et al., 2021). Thus, the question of what mediates ATG9A recruitment/tethering to ubiquitin-rich condensates is still open and should, at least in part, be addressed by defining specific ATG9A domains (or vesicle components) that bind to poly-ubiquitin, autophagy adapters, or other constituents of the ubiquitin-rich condensate.

Our live-cell imaging experiments, taken together with previous work (Broadbent et al., 2023), suggest that although autophagy foci rapidly assemble and disassemble in the absence of nutrient stress, stable assembly of autophagy foci to form committed autophagosomes can occur at ubiquitin-rich condensates. These observations support the model that ubiquitinated protein condensates recruit ATG proteins to stimulate autophagosome formation (Fujioka and Noda, 2021; Turco et al., 2021). Our work also in-part defined the hierarchy of protein recruitment to the ubiquitin-rich condensates, with ATG9A being upstream of ATG2 recruitment to the condensates. Perhaps most interestingly, it suggests a model in which autophagy machinery ‘samples’ the cellular environment until it senses the condensation of poly-ubiquitinated cargo, which then trigger autophagosome formation. It seems likely, based on this study and previous work (Broadbent et al., 2023; Olivas et al., 2023; van Vliet et al., 2022), that ATG9A recruitment to ubiquitin-rich condensates provides an early signal to recruit ATG2 and establish a lipid transfer complex between ER subdomains and the expanding ATG9A lipid compartment. Whether the ATG9A-ATG2 complex plays a direct role in recruiting other factors (e.g., the ULK1 complex) or whether they are recruited independently to ubiquitin-rich condensates remains unclear.

Our observations also suggest that deletion of the ATG9A-ATG2 lipid transfer complex affects p62 differently than loss of other ATG proteins (ATG5 and ATG13). In comparing these ATG KO lines, we did not detect a significant difference in the overall rate of p62 degradation, but instead deletion of either ATG2A/B or ATG9 resulted in a build-up of phosphorylated p62, which was not observed to the same degree in cells lacking LC3 conjugation or ULK1 complex activity (ATG5 KO and ATG13 KO, respectively). The phosphorylated form of p62 is thought to have higher affinity for ubiquitin and thus likely marks the pool of p62 within the ubiquitin-rich condensate (Ichimura et al., 2013; Matsumoto et al., 2015; Matsumoto et al., 2011; Pilli et al., 2012). Consistent with this hypothesis, we observed an increase in p62 puncta volume in ATG9A or ATG2A/B KO cell lines compared to ATG5 or ATG13 KO cells. The increase in p62 condensate size upon ATG9A/ATG2 deletion, without a change in p62 turnover rate compared to other ATG KOs, is intriguing. It suggests that the lipid transfer complex may play a role in regulating p62 condensate size. Indeed, it is possible that engagement of the lipid transfer complex at the condensate—a commitment step in autophagy initiation—inhibits condensate growth to ensure that condensate size does not exceed the capacity of the autophagosome (Agudo-Canalejo et al., 2021; Turco et al., 2021). Perhaps the ATG9A-ATG2 lipid transfer complex recruits a phosphatase to dephosphorylate p62 and inhibit its ability to recruit additional poly-ubiquitin into the condensate. Interestingly, the catalytic subunit of protein phosphatase-1 was identified in a proteomic analysis of ATG9A vesicles (Judith et al., 2019). Alternatively, individual autophagosomes could partially degrade condensates, eliminating them in a step wise process, rather than a single large autophagosome engulfing the entire condensate.

In summary, our work provides mechanistic insight into how the formation of p62-scaffolded ubiquitin-rich condensates trigger the recruitment of ATG9A to initiate autophagosome growth in the context of basal autophagy. Going forward, it will be critical to precisely define ATG9A interactions with components of ubiquitin-rich condensates and more broadly understand the role of these condensates as a platform for recruitment of other autophagy factors.

## Acknowledgements

We thank the Fritz B. Burns Foundation for generous student and instrument support. We thank the Simmons Center for Cancer Research for student fellowships to CMM and DMP. JLA is supported by a National Institutes of Health R01 from NIGMS (NIH R01 GM147310-01) and an American Cancer Society Research Scholar Grant (133550-RSG-19-006-01-CCG). JCS is supported by National Institute for General Medical Sciences (DP2 GM 142307). BCN and JCP are supported by National Institutes of Health R01 from NIA (R01AG066874).

## Materials and Methods

### Antibodies, chemicals, and reagents

| Antibodies | Source | Identifier |
| --- | --- | --- |
| Anti-SQSTM1/p62 mouse | Abcam | ab56416 |
| Anti-LC3B rabbit | Cell Signaling Technology | 3868S |
| Anti-actin mouse | Santa Cruz Biotechnology | sc-8432 |
| Anti-β-actin rabbit | Cell Signaling Technology | 4970S |
| Anti-phospho-SQSTM1/p62 (Thr269/Ser272) rabbit | Cell Signaling Technology | 13121S |
| Anti-phospho-SQSTM1/p62 (Ser349) rabbit | Cell Signaling Technology | 16177S |
| Anti-phospho-SQSTM1/p62 (Ser403) rabbit | Cell Signaling Technology | 39786S |
| Anti-Tax1bp1 rabbit | Cell Signaling Technology | 5105S |
| Anti-NBR1 rabbit | Cell Signaling Technology | 48046 (Product discontinued) |
| Anti-NDP52 rabbit | Cell Signaling Technology | 48046 (Product discontinued) |
| Anti-OPTN rabbit | Cell Signaling Technology | 48046 (Product discontinued) |
| Anti-HA-Tag mouse | Cell Signaling Technology | 2367S |
| Anti-HA-Tag goat | Novus Biologicals | NB600-362 |
| NBR1 | Cell Signaling Technology | D2E6 |
| NDP52 | Cell Signaling Technology | D1E4A |
| Anti-Ubiquitin rabbit | Cell Signaling Technology | 3933S (Product discontinued) |
| Anti-rabbit AlexaFluor 488 goat | Abcam | ab150077 |
| Anti-rabbit AlexaFluor 633 goat | Thermo Fisher Scientific | A-21071 |
| Anti-mouse AlexaFluor 488 goat | Thermo Fisher Scientific | A-11001 |
| Anti-mouse AlexaFluor 633 goat | Invitrogen | A-21126 |
| Anti-rabbit AlexaFluor 488 donkey | Thermo Fisher Scientific | A-21206 |
| Anti-mouse AlexaFluor 568 donkey | Thermo Fisher Scientific | A10037 |
| Anti-goat AlexaFluor 633 donkey | Thermo Fisher Scientific | A-21082 |

| Cells | Source | Identifier |
| --- | --- | --- |
| HEK293T | ATCC |  |
| HA-ATG9A KI HCT116 | Parent line from ATCC | - |
| U2OS, | ATCC HTB-96 |  |
| LentiX-293T | ATCC |  |

| Chemicals or Reagents | Source | Identifier |
| --- | --- | --- |
| Penicillin-Streptomycin | Gibco | 15140122 |
| Dulbecco's modified Eagle's medium | Gibco | 11965-092 |
| RPMI | Gibco | 11875093 |
| Fetal Bovine Serum | Genesee Scientific | 25-514 |
| Phosphate buffered saline (1X) | Corning | 21-040-CV |
| PEI-MAX | Polysciences | 24765-1 |
| Polybrene | Sigma-Aldrich | TR-1003-G |
| LICOR Intercept | LI-COR Biosciences | 927-70001 |
| Phosphoblocker™ Blocking Reagent | Cell Biolabs | AKR-103 |
| Puromycin Dihydrochloride | ThermoFisher Scientific | A1113802 |
| Wortmannin | Cell Signaling Technology | 9951S |
| Dimethyl Sulfoxide | Fisher Scientific | D136-1 |
| JFX650(HaloTag) | Grimm Et al. 2021 | N/A |
| JF503(SNAP Tag) | Grimm Et al. 2015 | N/A |
| 7-bromo heptonal (HaloTag Blocker) | ThermoFisher Scientific | H54762 |
| AP20187 | APEX BIO | B1274 |
| MLN7243 (TAK-243) | Cayman Chemical | 30108 |
| Paraformaldehyde | Sigma-Aldrich | 158127-100 |
| Triton X-100 | Fisher Scientific | BP151-500 |
| Tween-20 | Fisher Scientific | BP337-100 |
| SEA BLOCK Blocking Buffer | ThermoFisher Scientific | 37527 |
| DAPI | Cayman Chemical Company | 14285 |
| Prolong Diamond Antifade mounting reagent | Thermo Fisher Scientific | P36961 |

### Plasmids

| Plasmid | Source | Identifier |
| --- | --- | --- |
| pLenti-puro | Addgene | Plasmid #39481 |
| psPAX2 | Addgene | Plasmid #12260 |
| pMD2.G | Addgene | Plasmid #12259 |
| PX459 | Addgene | Plasmid #62988 |
| pBA-mCherry-EGFP-PIM | Addgene | Plasmid #111758 |
| AAVS1-Snap-P62-HRD |  |  |
| AAVS1-eGFP-LC3B-HRD |  |  |
| AAVS1 sgRNA |  |  |

### Primers

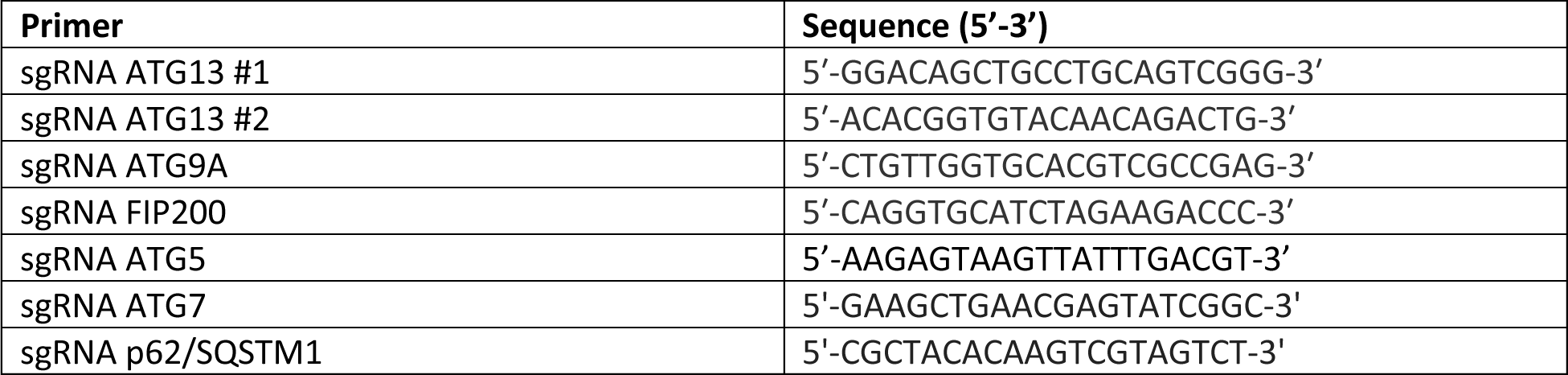

### Cell culture, transfection, and viral transduction

HEK293T cells, HCT116 cells, and their derivatives were cultured in Dulbecco’s modified Eagle’s medium (DMEM), supplemented with 10% fetal bovine serum (FBS) and Penicillin-Streptomycin (Pen-Strep) at 37°C in a 5% CO_2_ incubator. U2OS cells were cultured in RPMI media supplemented with FBS and Pen-Strep at 37°C in a 5% CO_2_. HEK293T and HCT116 cell were transiently transfected using polyethylenimine MAX (PEI-MAX) according to the manufacturer’s protocols. U2OS cells were transiently transfected with Lipofectamine 3000 or LONZA according to manufacturer’s protocols.

Lentivirus was generated using pLenti-puro (7 ug), psPAX2 (5.25 ug), and pMD2.G (1.75 ug) plasmids that were transfected in LentiX-293T cells within 15 cm plates with PEI-MAX at a 4:1 (ug:ug) ratio with cDNA. Virus was produced for 48 hours in the media and collected, centrifuged, and filtered through a sterile 0.45 um filter (Cat. No. SE1M003M00). The virus containing medium was then supplemented with 10 ug/mL of polybrene and allowed to incubate with the targeted cells for 24 hours at 37°C in a 5% CO_2_ incubator. The cells were washed and split for either chemical selection with puromycin or cell sorting.

### Generation of CRISPR-Cas9 knock-out and knock-in cell lines

To identify appropriate and effective sgRNAs, we utilized the CRISPR tool made by Concordet and Haeussler (http://crispor.tefor.net/) (Concordet and Haeussler, 2018). Using genomic exon DNA from the target proteins, we selected the highest scoring sgRNAs calculated by CRISPOR and ordered primers from Eton Bioscience, Inc. Using the cloning protocol defined by Zhang, et. al. (Cong et al., 2013), we cloned the primers into px330, px459 or px458. We then followed Lipofectamine 3000 or PEI-MAX transfection protocols and either chemically selected or cell sorted to enrich for CRISPR/Cas9-positive cells. After validating the efficiency of the sgRNAs in a mixed population of cells, we selected for single cell clones through serial dilution or single cell sorting into 96 well plates. The grown colonies were then tested by western blot to check for lack of protein expression. U2OS cells stably expressing HaloTag at their respective ATG loci are previously described and published (Broadbent et al., 2023). U2OS cells expressing Halo-ATG2A and an ATG9KO is also previously published (Broadbent et al., 2023). U2OS cells expressing Halo-ATG9A with ATG5KO, ATG2A/BKO, and ATG9KO are previously published (Perez et al., 2022).

### Immunoblotting

To prepare whole-cell protein lysates, cells were washed twice and harvested with ice-cold PBS. Cell pellets were resuspended in RIPA lysis buffer (25mM Tris-HCl [ph 7.5], 75 mM NaCl, 0.5% [wt/vol] Triton X-100, 2.5 mM EDTA, 0.05% [wt/vol] SDS, and 0.25% [wt/vol] Deoxycholate) supplemented with protease and phosphatase inhibitors and incubated for 15 min at 4°C with gentle rotation. The lysed cells were then centrifuged at 21,000xg for 10 min at 4°C and the supernatant collected. A protein assay is then performed to calculate then approximate protein concentration. Whenever a western blot was performed and whole cell lysates analyzed, 50 ug of protein was loaded into each well of the 4-5% Criterion^TM^ TGX^TM^ Precast Midi Protein Gel. After SDS-PAGE, the separated proteins were transferred onto a nitrocellulose membrane using an iBlot^TM^ 2. Membranes were allowed to dry completely and then soaked in PBS for 15-30 minutes and subsequently blocked with either LICOR Intercept blocking buffer or Phosphoblocker^TM^ for 1 hour at room temperature on a roller. Antibodies were diluted in blocking buffer according to the manufacturer’s specifications and allowed to incubate with the membrane in 4°C on a roller overnight. The membranes were washed once, 5 minutes each, with 0.1% Tween-20/PBS and twice with PBS and then LI-COR secondaries diluted 1:20,000 in PBS were incubated with the cells for 1 hour at room temperature on a roller. The membranes were washed again and imaged on the LI-COR Odyssey imaging system. Western blotting quantifications were calculated on Image Studio and statistical analyses were performed in GraphPad Prism 9. For U2OS protein lysates, cells were lysed in 300ul 1X sample buffer (BioRad) and boiled for 5 min at 95 C and loaded onto an or 4-20% TGX stain-free polyacrylamide gels (Biorad), followed by standard western blotting procedures. JFX650 was measured on ChemiDoc imaging system (BioRad) using the Cy5 filter set following by the stainfree loading control. Gels were transferred using the Trans-Blot Turbo system (with Turbo transfer buffer, BioRad) and blocked for 30 minutes in 5% milk. Primary antibody was left overnight at 1:3000 concentration. The blot was washed three times in .05% PBST and HRP-conjugated goat anti-mouse secondary was added at 1:2000 for 1 hour (Invitrogen, 31430, 1:2000). U2OS cell western blot quantification was carried out using the ImageQuant™ software (Cytiva).

### Deuterium Exchange

HEK293T WT and HEK293T ATG9A KO cells were seeded into 15cm plates at approximately 15% confluency and allowed to settle and grow overnight. A total of 15 plates per cell line were seeded, totaling 30 plates. The next day, DMEM/10% FBS cell culture medium with or without supplemented D_2_O (final D_2_O percentage at 5% to total volume) were added to the cells and allowed to incubate for 4, 12, 24, or 36 hours at 37°C. The cells containing 5% D_2_O cell culture medium were placed in a 5% CO_2_ incubator containing a 5% D_2_O water bath. The control cells (incubated in growth medium that did not contain added D_2_O) were incubated with a water bath containing no extra D_2_O. At each indicated time point, the cells were washed twice with ice cold PBS and collected in ice cold PBS via cell scraper into a previously weighed 15 mL conical tube. Pellets were centrifuged, supernatant aspirated, and centrifuged again to pull down excess supernatant. After aspiration of the excess supernatant, the tubes were weighed, and pellet size calculated. Cell lysates were generated by RIPA detergent lysis (milligram pellet:uL RIPA lysis buffer at a 1:10 ratio) and protein concentrations calculated for each sample. The proteins were then prepared for mass spectrometry as described below.

### Quantitative LC-MS/MS

After protein concentrations for each sample were calculated, 50 ug of protein from each sample was diluted in 6 M Guanidine/HCl and incubated for 5 min in 95°C. The samples were cooled and then placed in a VWR Centrifugal Filter with a 30 kD molecular weight cut-off (Cat. No. 82031-534) and centrifuged at 15,000xg for 15 minutes. The flow through was discarded and more 6 M Guanidine-HCl was added and washed through the filter by centrifugation at the same speed. 100 uL of 6 M Guanidine/HCl containing 9.2 mM of dithiothreitol (DTT) was added to the filter, vortexed briefly, and incubated at 60°C for 1 hour. After the samples were allowed to cool, iodoacetamide (IAM) was added for a final concentration of approximately 20 mM. The samples were protected from light and allowed to incubate at 25°C for 1 hour. The samples were then washed with 25 mM ammonium bicarbonate (ABC) three times, with centrifugation at 15,000xg for 15 minutes in between and the flow through discarded. The collection tubes were then washed with HPLC grade water three times. Mass spec grade Trypsin was diluted to 1ug Trypsin:100uL of 25 mM ABC (1 ug of Trypsin per 50 ug of protein) and added to each sample. The samples were mixed through mild vortexing, spun down briefly to pool the sample on top of the filter, and allowed to incubate on a shaker at 37°C for 18 hours. The samples were then centrifuged for 30 minutes at 15,000xg, with one more wash through of 25 mM ABC. The filtrant was collected and transferred to vials to be dried by speed-vacuum.

Mass spectrometry data were collected using an Orbitrap Fusion Lumos mass spectrometer (Thermo Fisher Scientific, Waltham, MA, USA) coupled to an EASY-nLC 1200 liquid chromatography (LC) pump (Thermo Fisher Scientific, Waltham, MA, USA). A capillary reverse-phase column (EASY-spray column pepMap ® RSLC, C18, 2 μm, 100 Å, 75 μm × 15 cm) was used for separation of peptides. The mobile phase was comprised of buffer A (0.1% formic acid in optima water) and buffer B (optima water and 0.1% formic acid in 80% acetonitrile). The peptides were eluted at 300 nL/min with the following gradients over 2 h: 3–25% B for 80 min; 25–35% B for 20 min; 35–45% B for 8 min; 45–85% B for 2 min and 85% for 8 min. Data were acquired using the top speed method (3 s cycle). A full scan MS at resolution of 120,000 at 200 m/z mass was acquired in the Orbitrap with a target value of 4e5 and a maximum injection time of 60 ms. Peptides with charge states of 2-4 were selected from the top abundant peaks by the quadrupole for high energy collisional dissociation (HCD with normalized energy 29) MS/MS, and the fragment ions were detected in the linear ion trap with target AGC value of 1e4 and a maximum injection time of 250 ms. The dynamic exclusion time was set at 40 s. Precursor ions with ambiguous charge states were not fragmented.

Kinetics data acquisitions were performed in MS-only mode and collected at 60,000 m/z resolution. These settings increase signal intensity, improve signal-to-noise, and give more scan points per elution chromatogram, greatly enhancing kinetic analysis accuracy (Naylor et al., 2017).

### Peptide identification and Protein Label Free Quantitation

PEAKS Studio software (version X) was used for de novo sequencing, and database searching to identify proteins in our raw MS data as well as to quantify, filter (quality-control), and normalize our quantitation data for each protein (Zhang et al., 2012). Peptides were identified from MS/MS spectra by searching against the Swiss-Prot human (2020) with a reverse sequence decoy database concatenated. Variables for the search were as follows: enzyme was set as trypsin with one missed cleavage site.

Carbamidomethylation of cysteine was set as a fixed modification while N-terminal acetylation and methionine oxidation were set as variable modifications. A false positive rate of 0.01 was required for peptides and proteins. Minimum length of peptide was set to 7 amino acids. At least 2 peptides were required for protein identification. The precursor mass error of 20 ppm was set for the precursor mass, and the mass error was set as 0.3 Da for the MSMS. Label-free quantitation was enabled with MS1 tolerance ±20ppm and a MS2 tolerance ±50 ppm, carbamidomethylation of cysteine was set as a fixed modification, while N-terminal acetylation and methionine oxidation were set as variable modifications. Peptide assignments with a false discovery rate less than 1% were included in comparative quantitative analyses and used to generate protein identification files for the quantitative and kinetic analyses. Protein quantitation was performed using the ‘Label Free Quantitation’ (LFQ) module in the PEAKS software package.

### Kinetic Protoemics analysis

Protein turnover rates were calculated using the DeuteRater software tool (Naylor et al., 2017). Briefly, data from the mass spectrometer was converted into the .mzML file format using the MSconvet tool (Kessner et al., 2008). Peak picking was used as the first filtering option, no other options were used. An ID file for DeuteRater to use was created using the output files of the PEAKS X Pro (Zhang et al., 2012) peptide identification software. The ID file, .mzML files, and the experimental data (time since the start of labeling for each sample and the final amount of deuterium present) were provided to DeuteRater.

DeuteRater calculates a turnover rate based on a rise to plateau fit of the changes in isotopic envelopes as deuterium is incorporated. Default DeuteRater settings were used.

To identify which proteins to include for analysis, a custom scoring system was created. First, peptides associated with each protein needed to be identified in at least 6 of the 15 total samples (15 for WT and 15 for ATG9A KO), meaning that there needs to be a high enough abundance of peptides identified in both the WT samples and ATG9A KO samples. Second, peptides associated with each protein need to be found in at least 4 of the 5 time points for WT and ATG9A KO, including the no deuterium control (control, 4 hours, 12 hours, 24 hours, and 36 hours). Third, the R^2^ value calculated for the slope generated from the combined rate graphs by Deuterator (Fig. 1D) must be above 0.5. Lastly, the calculated 95% confidence interval associated with a specific protein found in both the WT and ATG9A KO samples could not overlap. If a protein was found to follow these four criteria, then that protein was included in the analysis found in Figure 1B. All of the above calculations were performed in Microsoft Excel.

### Immunofluorescent microscopy

Cells were seeded onto acid-etched coverslips and allowed to culture for 24 hours before treatments and/or fixation. When the cells were ready for fixation, the cells were washed with ice-cold PBS three times and incubated with 4% paraformaldehyde (PFA)/PBS for 10 minutes at 37°C while protected from light. The cells were subsequently washed with ice-cold PBS three times and then permeabilized with 0.1% Triton X-100/PBS for 10 minutes at room temperature on a gentle rocker while protected from light. The cells were then washed three times with ice-cold PBS and then blocked with SEA BLOCK Blocking Buffer for 1 hour at room temperature on a gentle rocker while protected from light. The blocking buffer was then aspirated from the cells and fresh blocking buffer containing diluted primary antibodies (diluted according to the manufacturer’s protocols) was added to the cells. The cells were placed in 4°C on a gentle rocker and allowed to incubate overnight while protected from light. The cells were then washed with ice-cold 0.1% Tween-20/PBS three times, five minutes for each wash, and PBS containing diluted secondaries (diluted according to the manufacturer’s protocols) was added to the cells and incubated at room temperature for 1 hours on a gentle rocker while protected from light. The cells were subsequently washed three times with ice cold 0.1% Tween-20/PBS, five minutes for each wash, and either mounted to microscope slides with Prolong Diamond Antifade mounting reagent or incubated with DAPI (1.43 uM final concentration in PBS) for five minutes and then mounted to microscope slides. The slides are then cured overnight at room temperature while protected from light. Images were acquired on a LEICA TCS SP8 confocal microscope fitted with a HC PL APO 63x/1.40 Oil CS2 objective and a HyD detection system (Leica Microsystems).

### Quantification of immunofluorescent microscopy

To increase reproducibility, each microscopy experiment was seeded, fixed, and stained on the same day with the same diluted antibodies in blocking buffer or PBS. Furthermore, the images collected had the same laser power and image resolution by set. Each analyzed image was processed using Huygens Essential express deconvolution tool on standard settings. Pearson’s coefficient was calculated using colocalization analyzer tool found in Huygens Essential. Threshold intensity values was set at either 1% or 10% of the highest intensity value for each channel, depending on how high the background was. The puncta volume was calculated using the 3D analysis software tool found in LAS-X. The statistical calculations were performed in GraphPad Prism 9.

### In-gel and immunoblot assessment of p62 degradation in ATG9A WT and M33 mutant

ATG9AKO edited U2OS cells were transfected using lipofectamine 3000 with 1 ug of a safe harbor AAVS1 locus homology recombination donor containing Halo-ATG9A WT or M33 mutant (K321L, K322L, K323L, T419W) cDNA and 1 ug of AAVS1 targeted sgRNA (Maeda et al., 2020; Xi et al., 2015). Cells were then selected for using puromycin at 1ug/ml for two weeks and sorted on HaloTag florescence. Single cell clones were grown and single cell clones expressing similar amounts of ATG9A WT and M33 were chosen for further experiments. The construct knocked into the cells contains a tet inducible promotor and this was utilized in our experiment. 300,000 cells were plated into a 6-well plate with and without doxycycline at 2ug/ml. The following day the cells were labeled with JFX650 at 100nM for 15 minutes, washed, and lysed in 300ul 1X sample buffer (BioRad) and boiled for 5 min at 95 C and loaded onto an or 4-20% TGX stain-free polyacrylamide gels (Biorad), followed by standard western blotting procedures. JFX650 was measured on ChemiDoc imaging system (BioRad) using the Cy5 filter set following by the stainfree loading control. Gels were transferred using the Trans-Blot Turbo system (with Turbo transfer buffer, BioRad) and blocked for 30 minutes in 5% milk. Primary antibody was left overnight at 1:3000 concentration. The blot was washed three times in .05% PBST and HRP-conjugated goat anti-mouse secondary was added at 1:2000 for 1 hour (Invitrogen, 31430, 1:2000). U2OS cell western blot quantification was carried out using the ImageQuant™ software (Cytiva). Statistical differences were evaluated by two-tail *t*-test.

### Live cell microscopy

Two microscopes were used for live cell microscopy in this manuscript. The first is an Olympus microscope (IX83), with a cellTIRF illuminator with four laser lines (100 mW 405nm, 200 mW 488nm, 300 mW 561nm, and 140 mW 640 nm) and an X-Cite TURBO multi-wavelength LED illumination system (Excelitas Technologies). The microscope is equipped with an environmental chamber (cellVivo) to control humidity, temperature, and CO_2_ level, a 100x TIRF oil-immersion objective (Olympus UApo N, NA = 1.49) with compatible excitation and emission filters. Alternatively, we used an i3 spinning disc confocal microscope equipped with a CSU-W1 confocal spinning disk system (Yokogawa), four laser lines (100 mW 405nm, 150 mW 488nm, 175 mW, 160 mW 561nm, and 140 mW 638 nm), a ORCA-Quest qCMOS camera.

### Chase-Pulse experiment with UAE Inhibitor and analysis

Halo-ATG9A C1 cell line was transfected with the AAVS1 safe locus HRD containing SNAP-p62 alongside an sgRNA and selected with puromycin. After selection, 200,000 cells were grown on glass coverslips (170 ± 5 μm, Schott). Coverslips were cleaned with 1M KOH (1h in a sonicated water bath), rinsed with ddH_2_O, cleaned with 100% ethanol (1h in a sonicated water bath), and dried under N_2_ stream before assembly on a 35 mm diameter imaging dish using epoxy. The day after seeding the HaloTag was blocked using 7-bromo-heptanol at 10 uM for 10 minutes. The cells were rinsed twice with media and incubated in 2 ml media to deplete excess halo ligand for 5 minutes. Cells were then treated by adding 2 ml of media containing 100nM JFX650 Halo ligand and 100 nM JF503 SNAP ligand. Drugs treatment was added alongside the dye by adding DMSO or UAE inhibitor at a concentration of 10uM for 9-10 hours. Prior to imaging, we swapped the media for 5 minutes, maintaining the drug during the wash and then added fresh treated media for imaging. Live-cell imaging was performed on a Olympus TIRF microscope using the 640 nm laser in a Highly Inclined and Laminated Optical sheet (HILO) angle and the Andor iXon 897 Ultra camera was used for signal detection. Image analysis was performed in an unbiased manner using scripts from ImageJ and ICY. Images were first prepped for analysis using ImageJ. Frame 5 of each movie was extracted and a rolling background subtraction of 20 pixels was applied to the image. These images were then analyzed using custom built ICY protocol that identifies foci using a wavelet filter and gaussian fitting spot detector to identify ROIs and the integrated intensity was quantified across all 3 replicates. Statistical differences were evaluated by two-tail *t*-test.

### Autophagosome foci kinetics with UAE inhibitor

Halo-ATG13 cells were stably transfected with GFP-LC3. 250,000 Cells were then plated on 35mm dish with 1.5H glass (Cellvis, product #: D35-20-1.5H) and left overnight. The following day, cells were labeled with JFX650 at 100nM for 15 min and washed for 5 min in media. Cells were then treated with and without UAE inhibitor at 10uM for 2 hours. Live-cell imaging was performed in standard conditions of 5% CO2 and 37 C. (description of microscope) Images were acquired at 1 frame every second for 8 minutes. For image analysis feasibility, images were scaled down to 727X512 pixels, and analyzed using a previously established pipeline called K-FOCUS (Barnaba et al., 2023).

### Live-cell imaging of Halo-ATG2 and 6X-ubiquitin-BFP

Endogenously edited Halo-ATG2A cells expressing WT ATG9A and KO were transfected with HA-6x-ubiquitin-eBFP2 by LONZA electroporation using standard settings for U2OS cells and a homemade buffer of RPMI supplemented with 50mM bicarbonate. Importantly, ATG9A KO is more prone to forming large aggregates, likely due to having basal nucleation of p62, so any cells that had excessive aggregation was not imaged keeping the two conditions more similar. Live-cell imaging was performed in standard conditions of 5% CO2 and 37 C. (description of microscope) Images were acquired at 1 frame every second for 8 minutes. For image analysis feasibility, images were scaled down and analyzed using a previously established pipeline called K-FOCUS (Barnaba et al., 2023). Statistical differences were evaluated by two-tail *t*-test.

### OPTN SE mutant with Halo-ATG9A KR mutant addback

ATG9A KO edited U2OS cells were transfected using lipofectamine 3000 with 1 ug of a safe harbor AAVS1 locus homology recombination donor containing WT ATG9A or KR mutations (K851R & K838R) cDNA and 1 ug of AAVS1 targeted sgRNA. Cells were selected using puromycin at 1ug/ml for two weeks. For the experiment, the selected cells were transfected with lipofectamine 3000 with 1 ug/ml of 6x-ubiquitin-BFP and 1 ug/ml of OPTN SE. Images were acquired on the Olympus microscope at 1 frame per second over 30 seconds.

## Supplemental Material

**Supplemental Figure 1.**
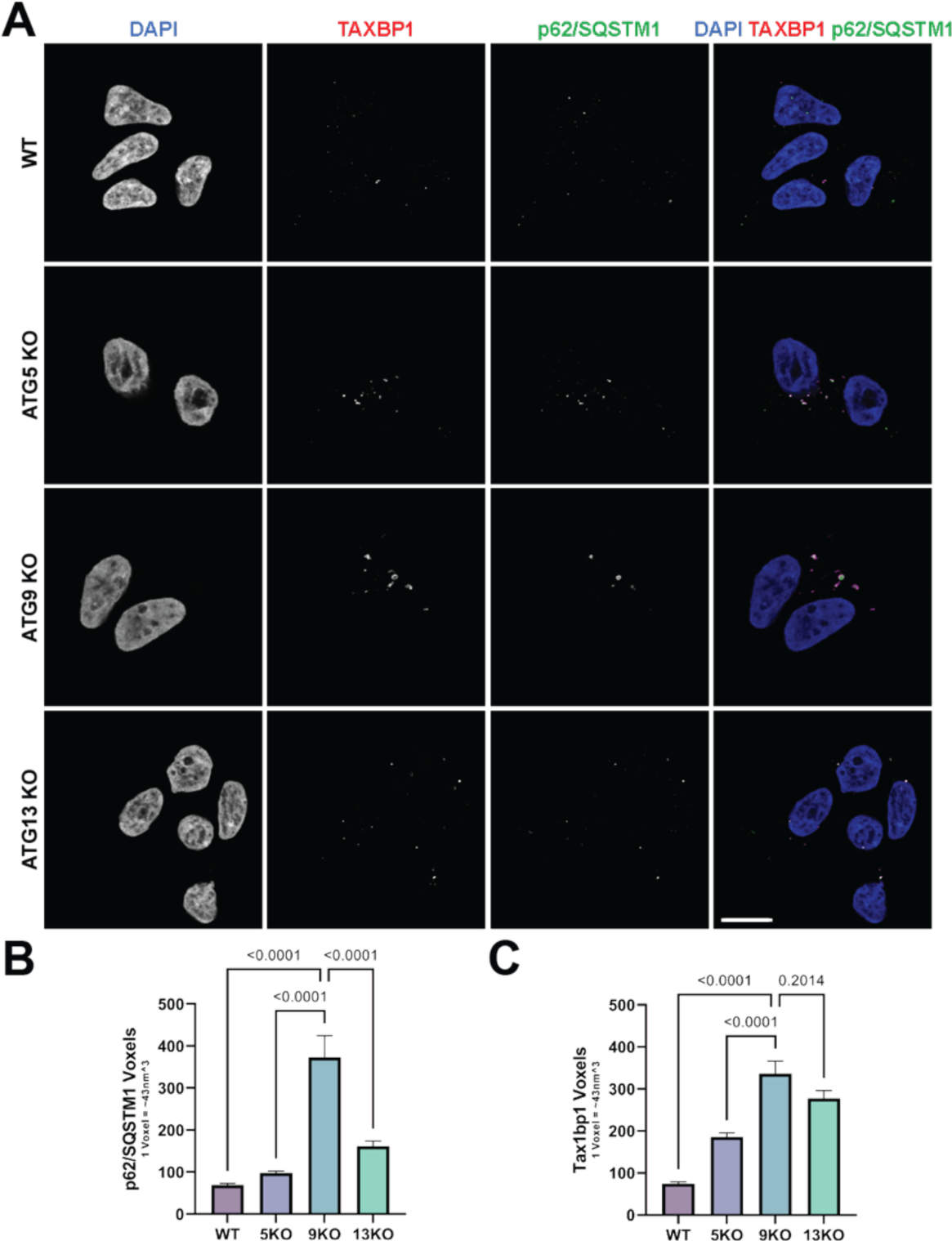
L**a**rge **p62 clusters in ATG KO cell lines colocalize with TAX1BP1 (A)** Confocal imaging of endogenous p62 and TAX1BP1 in WT and ATG5, ATG9A and ATG13 KO HCT-116 lines. **(B)** Quantification of endogenous p62 puncta size from the experiment in panel A. Quantitation is from 3 replicates with error bars representing SD. *P*-values were calculated using a two-tailed student *t*-test for pair-wise comparison. **(C)** Quantification of endogenous TAX1BP1 puncta size from the experiment in panel A. Quantitation is from 3 replicates with error bars representing SD. *P*-values were calculated using a two-tailed student *t*-test for pair-wise comparison

**Supplemental Figure 2.**
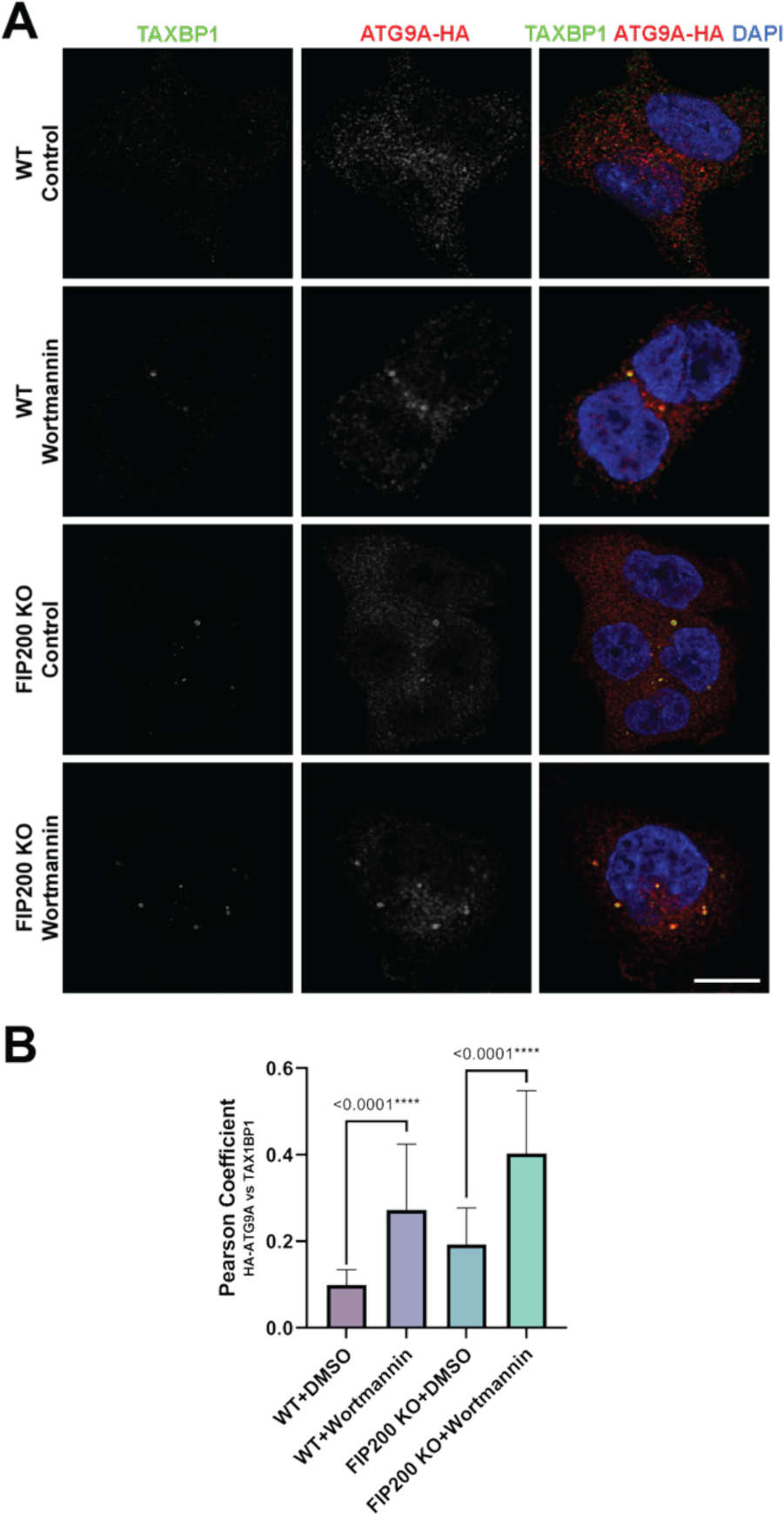
A**T**G9A **accumulates at p62 condensates in the absence of FIP200 (A)** Confocal imaging of endogenous TAX1BP1 and endogenously HA-tagged ATG9A in HCT-116 cells treated with or without 1 uM Wortmannin for 4 hours (scale bar = 10 µm) **(B)** Quantification of ATG9A-HA and FIP200 colocalization by Pearson’s coefficient. Quantitation is from 3 replicates with error bars representing SD. *P*-values were calculated using a two-tailed student *t*-test for pair-wise comparison

**Supplemental Figure 3.**
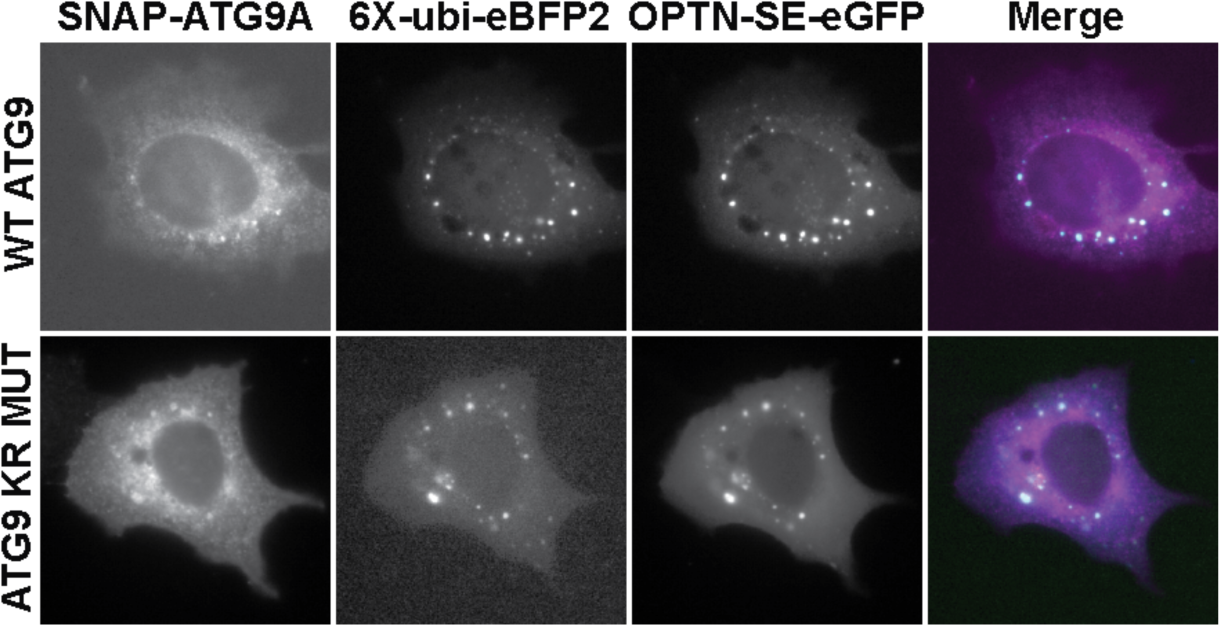
A**r**tificial **condensates composed of 6X-ubiquitin and OPTN recruit ATG9A.** U2OS cells stably expressing snap-ATG9A were transiently transfected with 6x-ubiquitin and mutant OPTN-SE. Imaging of SNAP, BFP2, and eGFP was done 1 frame/s.

## Supplemental Movie Legends

**Movie S1.** U2OS cells expressing Halo-ATG13 labeled with JFX650 (left panel, magenta in merge) and GFP-LC3B (center panel, green in merge) in control media imaged at 1 frame per second.

**Movie S2.** U2OS cells expressing Halo-ATG13 labeled with JFX650 (left panel, magenta in merge) and GFP-LC3B (center panel, green in merge) treated for 2 hours in media containing 10 µM MLN imaged at 1 frame per second.

**Movie S3.** U2OS cells expressing Halo-ATG9 labeled with JFX650 (left panel, magenta in merge) and SNAP-p62 labeled with JF503 (center panel, green in merge) in control media imaged at 1 frame per second.

**Movie S4.** U2OS cells expressing Halo-ATG9 labeled with JFX650 (left panel, magenta in merge) and SNAP-p62 labeled with JF503 (center panel, green in merge) after treatment with 10 µM MLN imaged at 1 frame per second.

**Movie S5.** U2OS cells expressing Halo-ATG2A labeled with JFX650 (left panel, magenta in merge) and 6x-Ubiquitin-BFP (center panel, green in merge) imaged at 1 frame per second.

**Movie S6.** ATG9A knock-out U2OS cells expressing Halo-ATG2A labeled with JFX650 (left panel, magenta in merge) and 6x-Ubiquitin-BFP (center panel, green in merge) imaged at 1 frame per second.

## REFERENCES

Adriaenssens, E., L. Ferrari, and S. Martens. 2022. Orchestration of selective autophagy by cargo receptors. Curr Biol. 32:R1357–R1371.

Agudo-Canalejo, J., S.W. Schultz, H. Chino, S.M. Migliano, C. Saito, I. Koyama-Honda, H. Stenmark, A. Brech, A.I. May, N. Mizushima, and R.L. Knorr. 2021. Wetting regulates autophagy of phase-separated compartments and the cytosol. Nature. 591:142–146.

Barnaba, C., D.G. Broadbent, G.I. Perez, and J.C. Schmidt. 2023. AMPK Regulates Phagophore-to-Autophagosome Maturation. bioRxiv:2023.2009.2028.559981.

Bjorkoy, G., T. Lamark, A. Brech, H. Outzen, M. Perander, A. Overvatn, H. Stenmark, and T. Johansen. 2005. p62/SQSTM1 forms protein aggregates degraded by autophagy and has a protective effect on huntingtin-induced cell death. J Cell Biol. 171:603–614.

Broadbent, D.G., C. Barnaba, G.I. Perez, and J.C. Schmidt. 2023. Quantitative analysis of autophagy reveals the role of ATG9 and ATG2 in autophagosome formation. J Cell Biol. 222.

Ciuffa, R., T. Lamark, A.K. Tarafder, A. Guesdon, S. Rybina, W.J. Hagen, T. Johansen, and C. Sachse. 2015. The selective autophagy receptor p62 forms a flexible filamentous helical scaffold. Cell Rep. 11:748–758.

Concordet, J.P., and M. Haeussler. 2018. CRISPOR: intuitive guide selection for CRISPR/Cas9 genome editing experiments and screens. Nucleic Acids Res. 46:W242–W245.

Cong, L., F.A. Ran, D. Cox, S. Lin, R. Barretto, N. Habib, P.D. Hsu, X. Wu, W. Jiang, L.A. Marraffini, and F. Zhang. 2013. Multiplex genome engineering using CRISPR/Cas systems. Science. 339:819–823.

Debnath, J., N. Gammoh, and K.M. Ryan. 2023. Autophagy and autophagy-related pathways in cancer. Nat Rev Mol Cell Biol. 24:560–575.

DeJesus, R., F. Moretti, G. McAllister, Z. Wang, P. Bergman, S. Liu, E. Frias, J. Alford, J.S. Reece-Hoyes, A. Lindeman, J. Kelliher, C. Russ, J. Knehr, W. Carbone, M. Beibel, G. Roma, A. Ng, J.A. Tallarico, J.A. Porter, R.J. Xavier, C. Mickanin, L.O. Murphy, G.R. Hoffman, and B. Nyfeler. 2016. Functional CRISPR screening identifies the ufmylation pathway as a regulator of SQSTM1/p62. Elife. 5.

Fujioka, Y., and N.N. Noda. 2021. Biomolecular condensates in autophagy regulation. Current opinion in cell biology. 69:23–29.

Ghanbarpour, A., D.P. Valverde, T.J. Melia, and K.M. Reinisch. 2021. A model for a partnership of lipid transfer proteins and scramblases in membrane expansion and organelle biogenesis. Proc Natl Acad Sci U S A. 118.

Ichimura, Y., S. Waguri, Y.S. Sou, S. Kageyama, J. Hasegawa, R. Ishimura, T. Saito, Y. Yang, T. Kouno, T. Fukutomi, T. Hoshii, A. Hirao, K. Takagi, T. Mizushima, H. Motohashi, M.S. Lee, T. Yoshimori, K. Tanaka, M. Yamamoto, and M. Komatsu. 2013. Phosphorylation of p62 activates the Keap1-Nrf2 pathway during selective autophagy. Mol Cell. 51:618–631.

Imai, K., F. Hao, N. Fujita, Y. Tsuji, Y. Oe, Y. Araki, M. Hamasaki, T. Noda, and T. Yoshimori. 2016. Atg9A trafficking through the recycling endosomes is required for autophagosome formation. Journal of cell science. 129:3781–3791.

Judith, D., H.B.J. Jefferies, S. Boeing, D. Frith, A.P. Snijders, and S.A. Tooze. 2019. ATG9A shapes the forming autophagosome through Arfaptin 2 and phosphatidylinositol 4-kinase IIIbeta. J Cell Biol. 218:1634–1652.

Kannangara, A.R., D.M. Poole, C.M. McEwan, J.C. Youngs, V.K. Weerasekara, A.M. Thornock, M.T. Lazaro, E.R. Balasooriya, L.M. Oh, E.J. Soderblom, J.J. Lee, D.L. Simmons, and J.L. Andersen. 2021. BioID reveals an ATG9A interaction with ATG13-ATG101 in the degradation of p62/SQSTM1-ubiquitin clusters. EMBO Rep:e51136.

Karanasios, E., S.A. Walker, H. Okkenhaug, M. Manifava, E. Hummel, H. Zimmermann, Q. Ahmed, M.C. Domart, L. Collinson, and N.T. Ktistakis. 2016. Autophagy initiation by ULK complex assembly on ER tubulovesicular regions marked by ATG9 vesicles. Nature communications. 7:12420.

Kessner, D., M. Chambers, R. Burke, D. Agus, and P. Mallick. 2008. ProteoWizard: open source software for rapid proteomics tools development. Bioinformatics. 24:2534–2536.

Kuma, A., M. Hatano, M. Matsui, A. Yamamoto, H. Nakaya, T. Yoshimori, Y. Ohsumi, T. Tokuhisa, and N. Mizushima. 2004. The role of autophagy during the early neonatal starvation period. Nature. 432:1032–1036.

Maeda, S., H. Yamamoto, L.N. Kinch, C.M. Garza, S. Takahashi, C. Otomo, N.V. Grishin, S. Forli, N. Mizushima, and T. Otomo. 2020. Structure, lipid scrambling activity and role in autophagosome formation of ATG9A. Nat Struct Mol Biol. 27:1194–1201.

Mari, M., J. Griffith, E. Rieter, L. Krishnappa, D.J. Klionsky, and F. Reggiori. 2010. An Atg9-containing compartment that functions in the early steps of autophagosome biogenesis. J Cell Biol. 190:1005–1022.

Mathis, A.D., B.C. Naylor, R.H. Carson, E. Evans, J. Harwell, J. Knecht, E. Hexem, F.F. Peelor, 3rd, B.F. Miller, K.L. Hamilton, M.K. Transtrum, B.T. Bikman, and J.C. Price. 2017. Mechanisms of In Vivo Ribosome Maintenance Change in Response to Nutrient Signals. Mol Cell Proteomics. 16:243-254.

Matoba, K., T. Kotani, A. Tsutsumi, T. Tsuji, T. Mori, D. Noshiro, Y. Sugita, N. Nomura, S. Iwata, Y. Ohsumi, T. Fujimoto, H. Nakatogawa, M. Kikkawa, and N.N. Noda. 2020. Atg9 is a lipid scramblase that mediates autophagosomal membrane expansion. Nat Struct Mol Biol. 27:1185–1193.

Matsumoto, G., T. Shimogori, N. Hattori, and N. Nukina. 2015. TBK1 controls autophagosomal engulfment of polyubiquitinated mitochondria through p62/SQSTM1 phosphorylation. Human molecular genetics. 24:4429–4442.

Matsumoto, G., K. Wada, M. Okuno, M. Kurosawa, and N. Nukina. 2011. Serine 403 phosphorylation of p62/SQSTM1 regulates selective autophagic clearance of ubiquitinated proteins. Mol Cell. 44:279–289.

Mizushima, N., and B. Levine. 2020. Autophagy in Human Diseases. N Engl J Med. 383:1564–1576.

Naylor, B.C., M.T. Porter, E. Wilson, A. Herring, S. Lofthouse, A. Hannemann, S.R. Piccolo, A.L. Rockwood, and J.C. Price. 2017. DeuteRater: a tool for quantifying peptide isotope precision and kinetic proteomics. Bioinformatics. 33:1514–1520.

Nishimura, T., N. Tamura, N. Kono, Y. Shimanaka, H. Arai, H. Yamamoto, and N. Mizushima. 2017. Autophagosome formation is initiated at phosphatidylinositol synthase-enriched ER subdomains. EMBO J. 36:1719–1735.

Olivas, T.J., Y. Wu, S. Yu, L. Luan, P. Choi, E.D. Guinn, S. Nag, P.V. De Camilli, K. Gupta, and T.J. Melia. 2023. ATG9 vesicles comprise the seed membrane of mammalian autophagosomes. J Cell Biol. 222.

Orsi, A., M. Razi, H.C. Dooley, D. Robinson, A.E. Weston, L.M. Collinson, and S.A. Tooze. 2012. Dynamic and transient interactions of Atg9 with autophagosomes, but not membrane integration, are required for autophagy. Mol Biol Cell. 23:1860–1873.

Pankiv, S., T.H. Clausen, T. Lamark, A. Brech, J.A. Bruun, H. Outzen, A. Overvatn, G. Bjorkoy, and T. Johansen. 2007. p62/SQSTM1 binds directly to Atg8/LC3 to facilitate degradation of ubiquitinated protein aggregates by autophagy. J Biol Chem. 282:24131–24145.

Perez, G.I., D. Broadbent, A.A. Zarea, B. Dolgikh, M.P. Bernard, A. Withrow, A. McGill, V. Toomajian, L.K. Thampy, J. Harkema, J.R. Walker, T.A. Kirkland, M.H. Bachmann, J. Schmidt, and M. Kanada. 2022. In Vitro and In Vivo Analysis of Extracellular Vesicle-Mediated Metastasis Using a Bright, Red-Shifted Bioluminescent Reporter Protein. Adv Genet (Hoboken). 3:2100055.

Pilli, M., J. Arko-Mensah, M. Ponpuak, E. Roberts, S. Master, M.A. Mandell, N. Dupont, W. Ornatowski, S. Jiang, S.B. Bradfute, J.A. Bruun, T.E. Hansen, T. Johansen, and V. Deretic. 2012. TBK-1 promotes autophagy-mediated antimicrobial defense by controlling autophagosome maturation. Immunity. 37:223–234.

Ravenhill, B.J., K.B. Boyle, N. von Muhlinen, C.J. Ellison, G.R. Masson, E.G. Otten, A. Foeglein, R. Williams, and F. Randow. 2019. The Cargo Receptor NDP52 Initiates Selective Autophagy by Recruiting the ULK Complex to Cytosol-Invading Bacteria. Mol Cell. 74:320–329 e326.

Ren, X., T.N. Nguyen, W.K. Lam, C.Z. Buffalo, M. Lazarou, A.L. Yokom, and J.H. Hurley. 2023. Structural basis for ATG9A recruitment to the ULK1 complex in mitophagy initiation. Sci Adv. 9:eadg2997.

Sawa-Makarska, J., V. Baumann, N. Coudevylle, S. von Bulow, V. Nogellova, C. Abert, M. Schuschnig, M. Graef, G. Hummer, and S. Martens. 2020. Reconstitution of autophagosome nucleation defines Atg9 vesicles as seeds for membrane formation. Science. 369.

Shoemaker, C.J., T.Q. Huang, N.R. Weir, N.J. Polyakov, S.W. Schultz, and V. Denic. 2019. CRISPR screening using an expanded toolkit of autophagy reporters identifies TMEM41B as a novel autophagy factor. PLoS Biol. 17:e2007044.

Sun, D., R. Wu, J. Zheng, P. Li, and L. Yu. 2018. Polyubiquitin chain-induced p62 phase separation drives autophagic cargo segregation. Cell Res. 28:405–415.

Turco, E., A. Savova, F. Gere, L. Ferrari, J. Romanov, M. Schuschnig, and S. Martens. 2021. Reconstitution defines the roles of p62, NBR1 and TAX1BP1 in ubiquitin condensate formation and autophagy initiation. Nature communications. 12:5212.

Turco, E., M. Witt, C. Abert, T. Bock-Bierbaum, M.Y. Su, R. Trapannone, M. Sztacho, A. Danieli, X. Shi, G. Zaffagnini, A. Gamper, M. Schuschnig, D. Fracchiolla, D. Bernklau, J. Romanov, M. Hartl, J.H. Hurley, O. Daumke, and S. Martens. 2019. FIP200 Claw Domain Binding to p62 Promotes Autophagosome Formation at Ubiquitin Condensates. Mol Cell. 74:330–346 e311.

van Vliet, A.R., G.N. Chiduza, S.L. Maslen, V.E. Pye, D. Joshi, S. De Tito, H.B.J. Jefferies, E. Christodoulou, C. Roustan, E. Punch, J.H. Hervas, N. O’Reilly, J.M. Skehel, P. Cherepanov, and S.A. Tooze. 2022. ATG9A and ATG2A form a heteromeric complex essential for autophagosome formation. Mol Cell. 82:4324–4339 e4328.

Vargas, J.N.S., M. Hamasaki, T. Kawabata, R.J. Youle, and T. Yoshimori. 2023. The mechanisms and roles of selective autophagy in mammals. Nat Rev Mol Cell Biol. 24:167–185.

Vargas, J.N.S., C. Wang, E. Bunker, L. Hao, D. Maric, G. Schiavo, F. Randow, and R.J. Youle. 2019. Spatiotemporal Control of ULK1 Activation by NDP52 and TBK1 during Selective Autophagy. Mol Cell. 74:347–362 e346.

Wang, Y.T., T.Y. Liu, C.H. Shen, S.Y. Lin, C.C. Hung, L.C. Hsu, and G.C. Chen. 2022. K48/K63-linked polyubiquitination of ATG9A by TRAF6 E3 ligase regulates oxidative stress-induced autophagy. Cell Rep. 38:110354.

Wurzer, B., G. Zaffagnini, D. Fracchiolla, E. Turco, C. Abert, J. Romanov, and S. Martens. 2015. Oligomerization of p62 allows for selection of ubiquitinated cargo and isolation membrane during selective autophagy. Elife. 4:e08941.

Xi, L., J.C. Schmidt, A.J. Zaug, D.R. Ascarrunz, and T.R. Cech. 2015. A novel two-step genome editing strategy with CRISPR-Cas9 provides new insights into telomerase action and TERT gene expression. Genome Biol. 16:231.

Yamamoto, H., S. Kakuta, T.M. Watanabe, A. Kitamura, T. Sekito, C. Kondo-Kakuta, R. Ichikawa, M. Kinjo, and Y. Ohsumi. 2012. Atg9 vesicles are an important membrane source during early steps of autophagosome formation. J Cell Biol. 198:219–233.

Yamano, K., R. Kikuchi, W. Kojima, R. Hayashida, F. Koyano, J. Kawawaki, T. Shoda, Y. Demizu, M. Naito, K. Tanaka, and N. Matsuda. 2020. Critical role of mitochondrial ubiquitination and the OPTN-ATG9A axis in mitophagy. J Cell Biol. 219.

Young, A.R., E.Y. Chan, X.W. Hu, R. Kochl, S.G. Crawshaw, S. High, D.W. Hailey, J. Lippincott-Schwartz, and S.A. Tooze. 2006. Starvation and ULK1-dependent cycling of mammalian Atg9 between the TGN and endosomes. Journal of cell science. 119:3888–3900.

Zachari, M., S.R. Gudmundsson, Z. Li, M. Manifava, R. Shah, M. Smith, J. Stronge, E. Karanasios, C. Piunti, C. Kishi-Itakura, H. Vihinen, E. Jokitalo, J.L. Guan, F. Buss, A.M. Smith, S.A. Walker, E.L. Eskelinen, and N.T. Ktistakis. 2019. Selective Autophagy of Mitochondria on a Ubiquitin-Endoplasmic-Reticulum Platform. Dev Cell. 50:627–643 e625.

Zaffagnini, G., A. Savova, A. Danieli, J. Romanov, S. Tremel, M. Ebner, T. Peterbauer, M. Sztacho, R. Trapannone, A.K. Tarafder, C. Sachse, and S. Martens. 2018. p62 filaments capture and present ubiquitinated cargos for autophagy. EMBO J. 37.

Zhang, J., L. Xin, B. Shan, W. Chen, M. Xie, D. Yuen, W. Zhang, Z. Zhang, G.A. Lajoie, and B. Ma. 2012. PEAKS DB: de novo sequencing assisted database search for sensitive and accurate peptide identification. Mol Cell Proteomics. 11:M111 010587.

